# Toxicity of chlordane at early developmental stage of zebrafish

**DOI:** 10.1101/119248

**Authors:** Jingxuan Xiong

## Abstract

Chlordane is highly toxic organochlorine pesticides that have been widely used throughout the world for decades and posing adverse effects on the environment. Contents detected in tissue and blood samples have resulted in a raising concern for their potential effects on wildlife and humans. In this study, we investigate the potential effect of chlordane on the development of zebrafish embryos. Zebrafish larvae were treated with different concentrations (0, 25, 50, 100, 200 ng/L) of chlordane from 12 hours postfertilization (hpf). Different early stage parameters were observed at 1, 2, 3 and 4 day post-fertilization (dpf). Chlordane-exposed zebrafish larvae appeared significant lower survival rate, developmental and hatching time delay and decreased embryo productivity. The heartbeat rate and blood flow were decreased in a dose dependent manner. These results suggested that exposure to real life of chlordane led to direct morphological and phenotypic changes and effects systems related to development and reproduction even in short-term manner.

## Introduction

Chlordane, belongs to organochlorine pesticides, is widely distributed contaminant in environments. Because of its high toxicity, stability, liposolubility and long biological halflives, chlordane has been considered one of the most-dangerous pesticides. Organochlorine pesticides, which have carcinogenic, teratogenic and endocrine-disruptive effects in animals [1-4], can maintain high concentrations of biomagnification and bioaccumulation within food chains [5-9]. Chlordane has been shown to have multi and alterable biological functions including estrogen-like functions [10-12], thus has detrimental effects on animal reproductive systems [13-15]. Chlordane has been detected consistently in many animal ecosystems even though it has been prohibited to use for decades in world-widely range [16-18]. Therefore, the impact of chlordane residues on the health of wildlife and humans has become an intractable concern.

Previous studies have reported that xenoestrogen treatment changes hormone levels and reproductive behaviors [19-21]. Zebrafish are frequently used as test animals in aquatic systems to examine the effect of endocrine-disrupting chemicals [22-25]. In zebrafish, the biological roles of the vertebrate-similar steroid hormones, testosterone and estrogen have been explored [26-28]. Expose to vertebrate estrogen can result in maturation in fish [29-31], which also affect the sexual development and sexual hormone regulation [32-35].

Zebrafish are well suitable for phenotypic screening and chemical toxicity [36-38]. The transparency of zebrafish embryos and high conservative evolutionary relationship to vertebrates, which make zebrafish a suitable model to investigate the genetic basis of human diseases [39-41]. The forward genetic screens of zebrafish makes this model powerful for discovery of novel gene functions in physiological process [42, 43]. Knockdown and knockout methodologies have been successfully applied in zebrafish [44, 45]. The short life span of small laboratory animals such as zebrafish represents an important advantage model to test programs in the environmental risk assessment of chemical toxicities [46].

In this study, we assessed the potential risk associated with the presence of chlordane in the aquatic environment by examining the morphological abnormalities, survival, hatching and heartbeat rates of zebrafish treated with chlordane.

## Materials and Methods

### Chemical

Technical-grade chlordane was purchased from Sigma-Aldrich (Cat. No: 45378-250MG, USA.). Stock solution was prepared by dissolving 100 mg chlordane in 10 mL water and stored at -20 °C. Four selected concentrations were prepared by dilution of the stock solution with final concentration of E3 medium (5 mM NaCl, 0.17 mM KCl, 0.33 mM CaCl and 0.33 mM MgSO_4_, PH 7.4).

### Zebrafish maintenance

Wild type AB zebrafish (*Danio rerio*) were at 28.5 °C on a 14-h-light/10-h-dark cycle. Embryos were obtained from the natural spawning of adult fish set up in pairwise crosses. All studies were performed with approval of the Jianghan University Animal Care and Use Committee (IACUC), protocol number 16-051. Embryos were incubated in E3 medium and staged according to previous literature [47].

### Chlordane treatment

Two hundred embryos of the same developmental stage were collected and place in 100 mL E3 medium and treated with four concentrations of chlordane (25, 50, 100, 200 ng/L) from 12 hpf. Non-treated embryos were set as controls maintained in the same medium without chlordane. These embryos were incubated at 28.5 °C and fresh medium were exchanged twice daily. Embryos were examined for survival rates at 1, 2, 3 and 4 dpf, and dead embryos were discarded. All the experiments were repeated in three replicates.

### Microscopy observation

Survival rates were calculated by counting the living larvae to the total embryos. Dead larvae were judged via the appearance of blood circulation, heartbeat and body color changes. Dead embryos were counted and discarded daily. The heartbeat rates of larvae from each group at corresponding time points were observed under a microscope and recorded (beats/minute) by using a timer. The hatching rates and morphological abnormalities were examined and observed daily.

### Statistical analysis

Statistical analysis was performed using SPSS (IBM, Armonk, NY). Graphs show mean and standard error of the mean unless otherwise specified. Significance for normally distributed data sets was calculated using the two-tailed Student t-test. *P* values < 0.05 were considered significant.

## Results

### Exposed to chlordane caused increasing mortality and abnormalities in zebrafish

To determine the toxicity of chlordane exposure to zebrafish, we treated zebrafish embryos with 0, 25, 50, 100, 200 ng/L chlordane from 12 hpf. The survival rates were decreased with an increase in high concentrations of chlordane during developmental processes (Fig. 1). In non-treated group, we observed no overt mortality through the whole experimental period. At 2 and 3 dpf, chlordane induced significant mortality form a concentration of 100 ng/L whereas at 4 dpf, the mortality was induced significantly from a concentration of 200 ng/L. zebrafish embryos developed normally in control group. In the chlordane-exposed groups, we observed prominent morphological abnormalities listed in Table 1, which showed a concentration-dependent manner. At higher concentration of 200ng/L, the larvae did not response to touch and the blood flow was lower compared to other groups at the same point. These results suggested that the lowest toxic concentration of chlordane was 50 ng/L from 2 to 3 dpf zebrafish larvae.

**Table 1.**
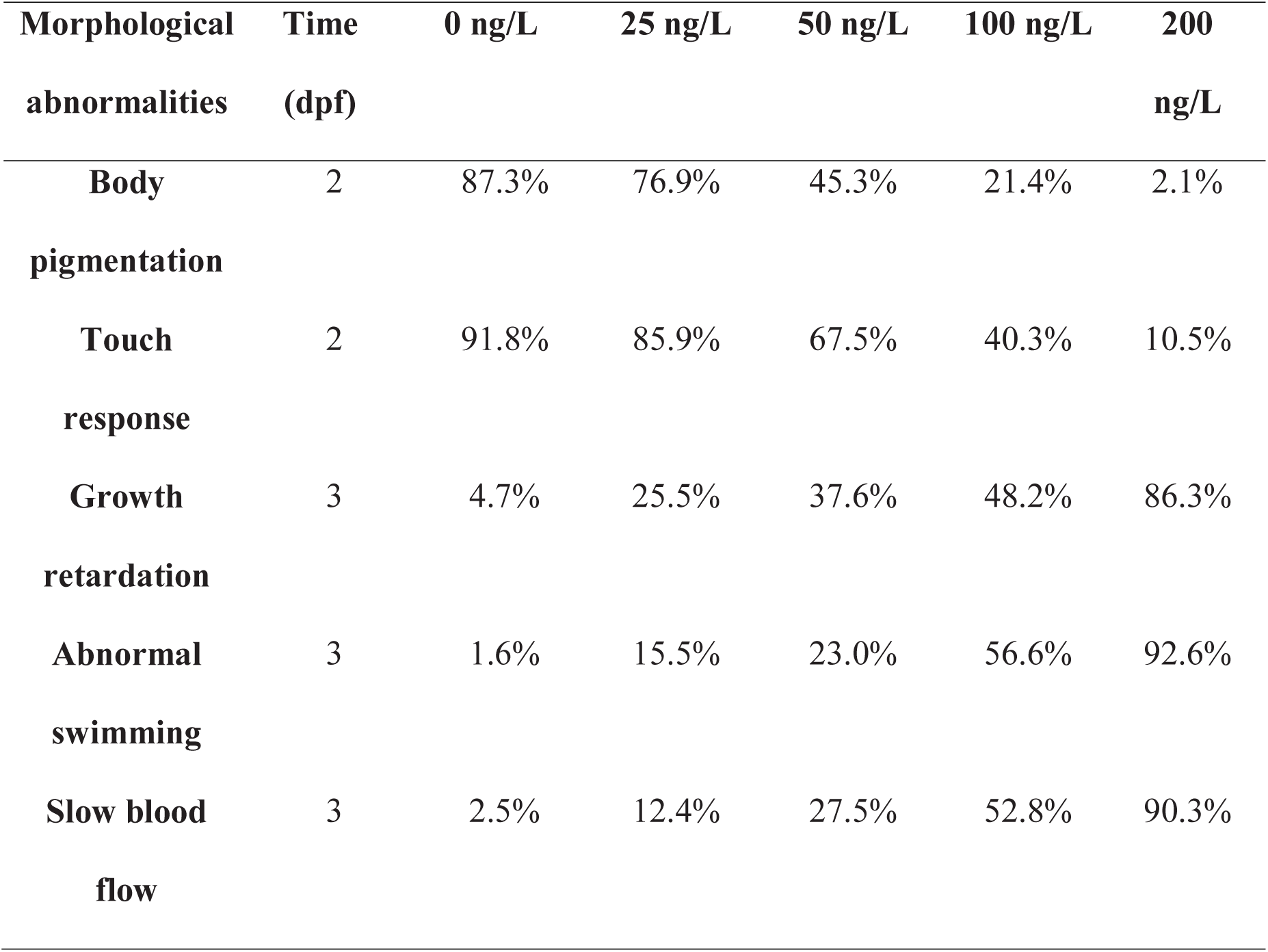
Effect of chlordane on zebrafish embryogenesis at different concentrations.

| **Morphological abnormalities** | **Time (dpf)** | **0 ng/L** | **25 ng/L** | **50 ng/L** | **100 ng/L** | **200 ng/L** |
| --- | --- | --- | --- | --- | --- | --- |
| **Body pigmentation** | 2 | 87.3% | 76.9% | 45.3% | 21.4% | 2.1% |
| **Touch response** | 2 | 91.8% | 85.9% | 67.5% | 40.3% | 10.5% |
| **Growth retardation** | 3 | 4.7% | 25.5% | 37.6% | 48.2% | 86.3% |
| **Abnormal swimming** | 3 | 1.6% | 15.5% | 23.0% | 56.6% | 92.6% |
| **Slow blood flow** | 3 | 2.5% | 12.4% | 27.5% | 52.8% | 90.3% |

**Figure 1.**
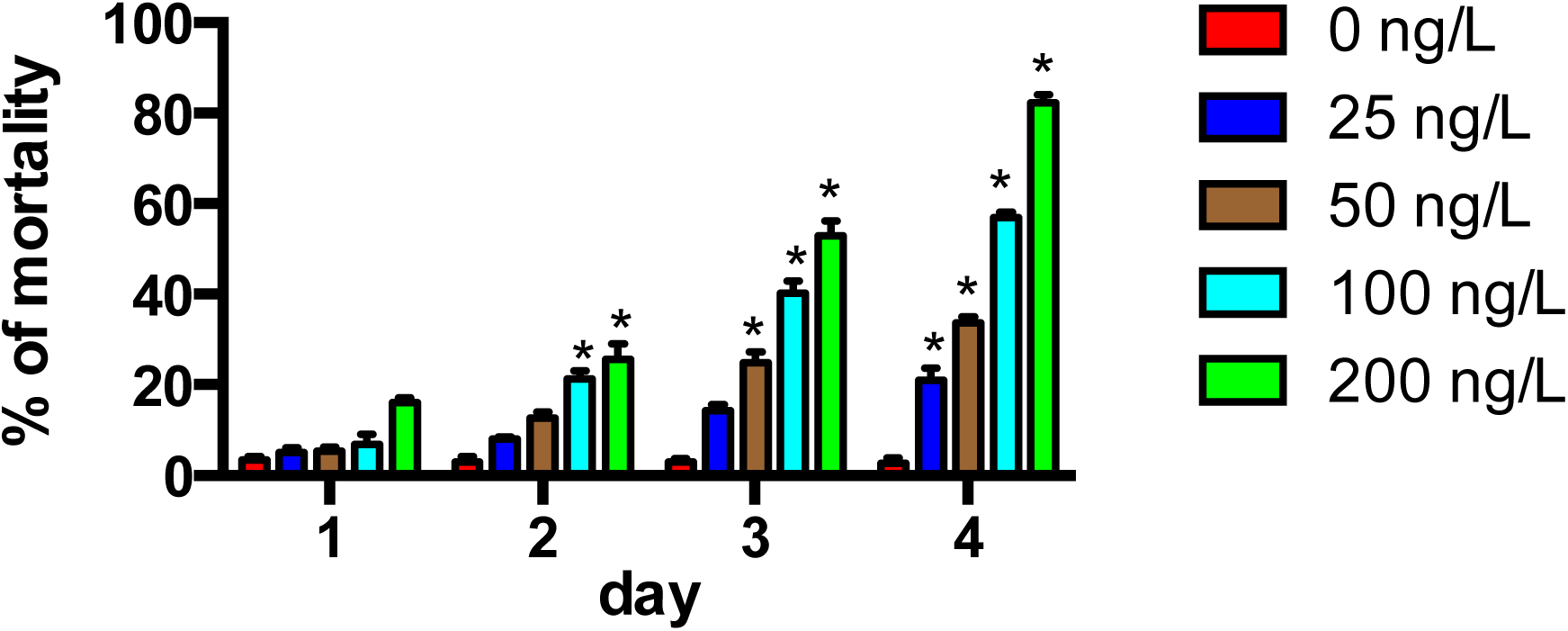
Survival rates in zebrafish-treated with different concentration of chlordane.

### Lower hatching rate in chlordane-treated zebrafish embryos

In the non-treated controls, zebrafish embryos were hatched sporadically between 2 and 3 dpf, however, the hatching rates were lower in the chlordane-exposed groups and had no hatching at higher chlordane concentration (200 ng/L) (Figure 2). The hatching rate in control group was 20.2% and 45% at 2 and 3 dpf, respectively, and 96.2% of embryos were hatched at 4 dpf. The similar percentage of hatching rates was observed at 25 ng/L chlordane-treated embryos. At concentrations of 50, 100 and 200 ng/L chlordane, zebrafish embryos started to hatch from 3 dpf and the percentage of hatching were significant lower compared to controls. The hatching rates of 50, 100 and 200 ng/L chlordane-treated groups were 40.6%, 15.1% and 5% at 4 dpf. The overall hatching rates in the embryos treated with chlordane were significantly decreased and the hatching time was obviously delayed.

**Figure 2.**
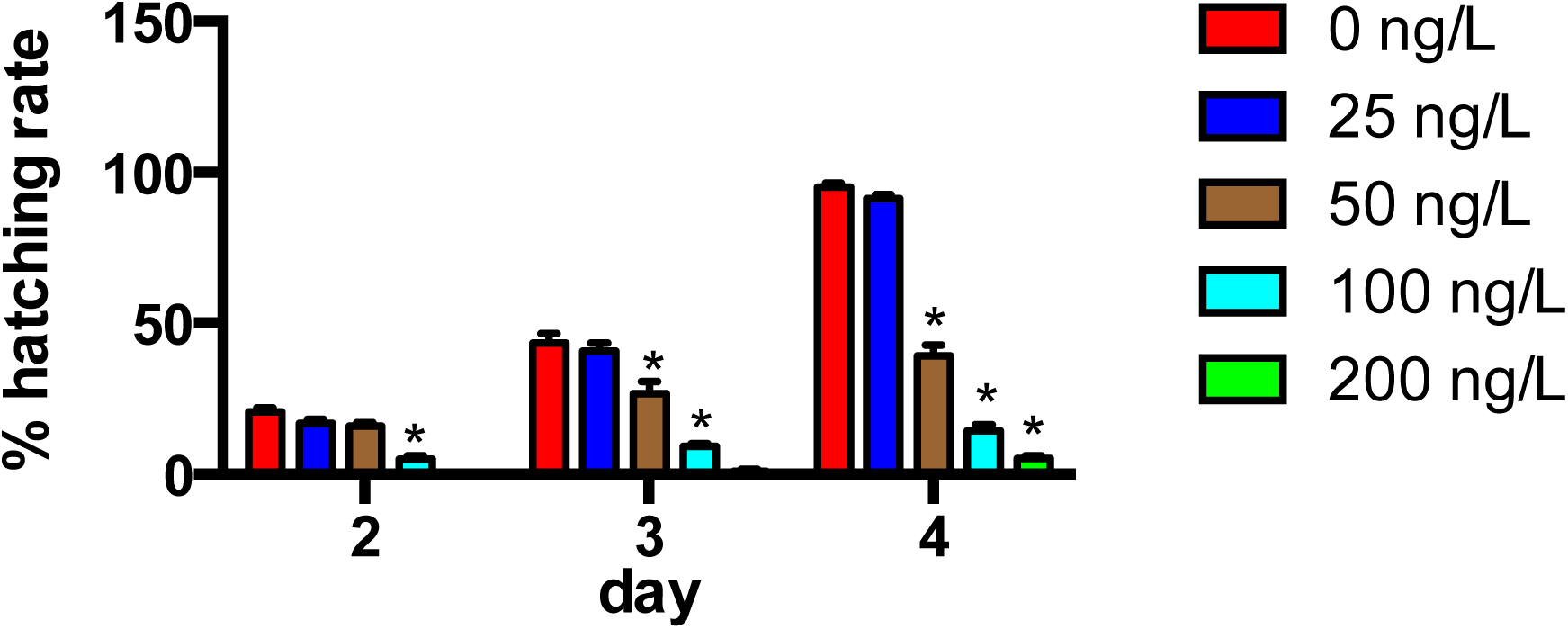
Hatching rates in zebrafish-exposed to different concentration of chlordane.

### Decreased heartbeat rates in chlordane-exposed zebrafish larvae

The heartbeat rates in the different chlordane exposure groups were monitored at the four developmental stages (Fig. 3). In the control group, heart rates increased and became prominent during development. We found that the number of beats/minute in controls was from 120/min to 140/min at 2 and 4 dpf, respectively. The heartbeat rats were significantly decreased in the chlordane-treated embryos and appeared a dose-dependent manner. In the 2 dpf, the number of beats/minute reduced from 110/min to 50/min in the 50 and 100 ng/L chlordane-treated embryos, respectively. In the 4 dpf, the beats decreased from 120/min to 30/min in the 50 and 100 ng/L chlordane-exposed groups, respectively. Observation made at 4 dpf has also shown the same trend until death was noticed. The heart beat rates were significantly decreased in all developmental stages at the four concentrations of chlordane compared to control zebrafish larvae.

**Figure 3.**
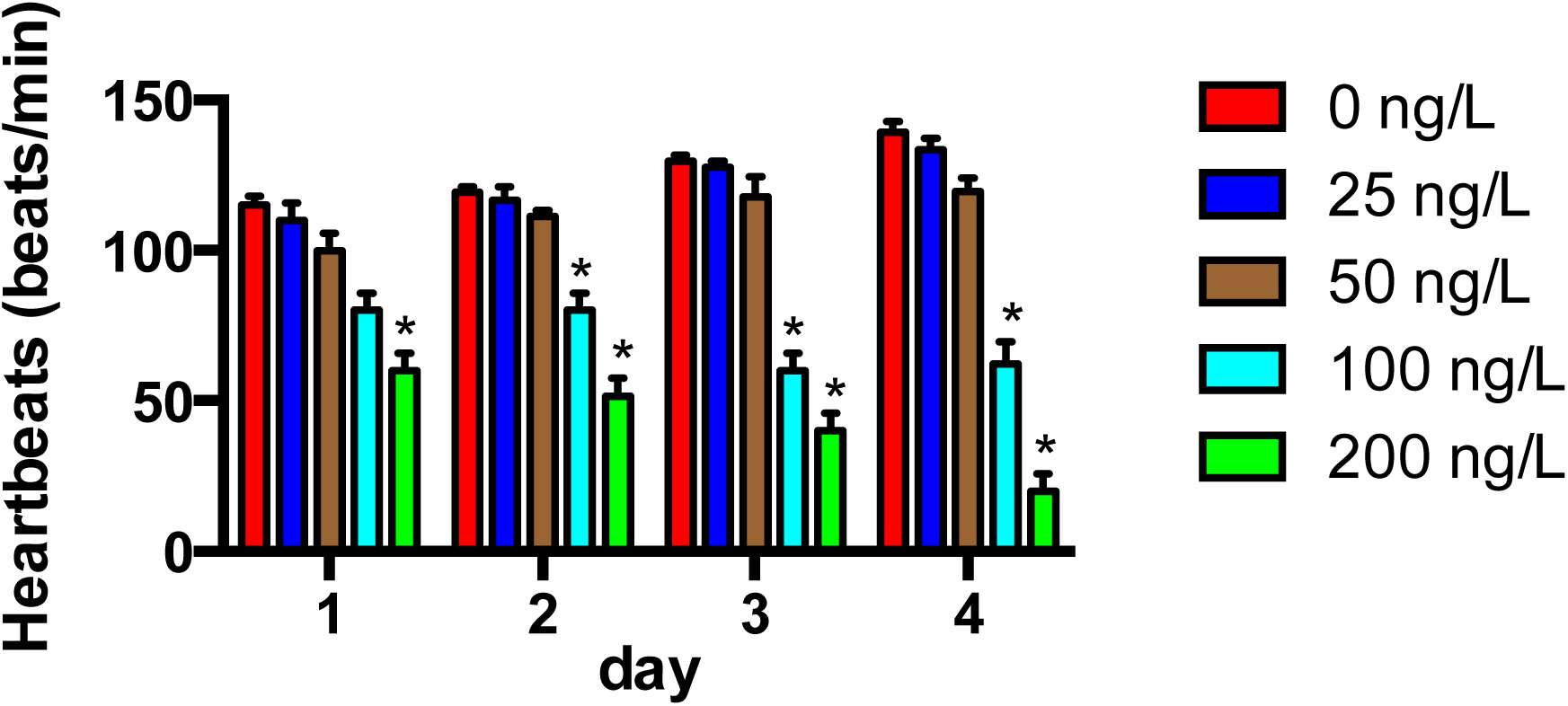
Heartbeat rates in zebrafish exposed to chlordane.

## Discussion

Numbers of pollutants can specifically affect the physiological processes occur within cells, tissues and organs during the embryogenesis and development [48-53]. In this study, we investigated the effects of chlordane on embryogenesis at four different time points during the zebrafish developmental stages from 2 to 4 dpf.

Early stage zebrafish embryos have been widely used for evaluating the toxicity and teratogenicity of chemicals might lead to severe impact on environmental and animal health [38, 54-58]. In the present study, we used four different concentrations of chlordane to examine the dose-dependent manner with developmental stages. These assays are useful for evaluating potential impacts on growth, survival and development of animals in the polluted environment and which can be used as valuable tools for environmental monitoring.

Chlordane led to a concentration-dependent manner increased in mortality (Fig. 1). The mortality was less than 3% in the controls during the test time period. Embryos (before 24 hpf) were more resistant than the larvae (2-4 dpf) when exposed to chlordane. The egg chorion may act a protective role to the embryos in a toxic environment. The lower hatching rates in the higher chlordane treated groups may due to the incorporation of chlordane into the embryos.

Hatching is an important point in the life cycle of fish and is well known as a crucial event during the embryogenesis [59-61]. Hatching rate delay in zebrafish embryos can happen when various reasons such as inhibition of the enzyme activities involved in hatching, which further result in behavioral defects and abnormal muscular movement. In the present study, we observed dose-dependent delay in the percentage of hatching rates (Fig. 2), which may be caused by these factors. Our data suggested that lower concentration of chlordane had little effect on hatching rates. However, further studies need be performed to investigate the role of chlordane on the enzymes activities involved in hatching process.

The heart rate is a key toxicology end point in the zebrafish embryonic test [56, 59, 62-64], which makes the heartbeat assessment as an important parameter in evaluating cardiac function. The number of beats/minute was slower in the chlordane-exposed larvae compared to the non-treated controls (Fig. 3). This effect might be possibly caused by malformation of the heart [65-67]. A weak heart resulted in defective functions and improper pumping of blood, which further led to retardation of body growth in larvae and slow blood flow (Table 1). These abnormalities in the chlordane-treated larvae indicated that chlordane was a cardiotoxic chemical. Thus, cardiotoxicity of chlordane might be the reason for abnormal morphological in this study, which further caused the larvae more lethargic at higher concentrations.

## Conclusion

Our study provided a good starting point for assessment of molecular mechanisms involved in development and embryogenesis that are impacted by chlordane. Further investigations will help to explore the potential mechanisms for pathogenesis as a consequence of chlordane exposure.

## References

1. Köhler, H.-R. and R. Triebskorn, Wildlife ecotoxicology of pesticides: can we track effects to the population level and beyond? Science, 2013. 341(6147): p. 759–765.

2. Cocco, P., On the rumors about the silent spring: review of the scientific evidence linking occupational and environmental pesticide exposure to endocrine disruption health effects. Cadernos de Saúde pública, 2002. 18(2): p. 379–402.

3. Balabanič, D., M. Rupnik, and A.K. Klemenčič, Negative impact of endocrine-disrupting compounds on human reproductive health. Reproduction, Fertility and Development, 2011. 23(3): p. 403–416.

4. McKinlay, R., et al., Endocrine disrupting pesticides: implications for risk assessment. Environment international, 2008. 34(2): p. 168–183.

5. Kutz, F.W., P.H. Wood, and D.P. Bottimore, Organochlorine pesticides and polychlorinated biphenyls in human adipose tissue, in Reviews of environmental contamination and toxicology. 1991, Springer. p. 1–82.

6. Hargrave, B., et al., Organochlorine pesticides and polychlorinated biphenyls in the Arctic Ocean food web. Archives of Environmental Contamination and Toxicology, 1992. 22(1): p. 41–54.

7. Corsolini, S., et al., Occurrence of organochlorine pesticides (OCPs) and their enantiomeric signatures, and concentrations of polybrominated diphenyl ethers (PBDEs) in the Adélie penguin food web, Antarctica. Environmental Pollution, 2006. 140(2): p. 371–382.

8. Edwards, C., Environmental pollution by pesticides. Vol. 3. 2013: Springer Science & Business Media.

9. Perry, A.S., et al., Insecticides in agriculture and environment: retrospects and prospects. 2013: Springer Science & Business Media.

10. Chighizola, C. and P.L. Meroni, The role of environmental estrogens and autoimmunity. Autoimmunity reviews, 2012. 11(6): p. A493–A501.

11. Rollerova, E. and M. Urbancikova, Intracellular estrogen receptors, their characterization and function (Review). Endocrine regulations, 2000. 34: p. 203-218.

12. McLachlan, J.A. and S.F. Arnold, Environmental estrogens. American Scientist, 1996. 84(5): p. 452–461.

13. Chapin, R., et al., The effects of perinatal/juvenile methoxychlor exposure on adult rat nervous, immune, and reproductive system function. Toxicological Sciences, 1997. 40(1): p. 138–157.

14. Fry, D.M., Reproductive effects in birds exposed to pesticides and industrial chemicals. Environmental Health Perspectives, 1995. 103(Suppl 7): p. 165.

15. Persson, S. and U. Magnusson, Environmental pollutants and alterations in the reproductive system in wild male mink (Neovison vison) from Sweden. Chemosphere, 2015. 120: p. 237–245.

16. Barni, M.F.S., et al., Persistent organic pollutants (POPs) in fish with different feeding habits inhabiting a shallow lake ecosystem. Science of the Total Environment, 2016. 550: p. 900–909.

17. Sanborn, J.R., et al., The fate of chlordane and toxaphene in a terrestrial-aquatic model ecosystem. Environmental Entomology, 1976. 5(3): p. 533–538.

18. Carriger, J.F., J. Castro, and G.M. Rand, Screening historical water quality monitoring data for chemicals of potential ecological concern: hazard assessment for selected inflow and outflow monitoring stations at the water conservation areas, South Florida. Water, Air, & Soil Pollution, 2016. 227(1): p. 27.

19. Vom Saal, F.S., et al., A physiologically based approach to the study of bisphenol A and other estrogenic chemicals on the size of reproductive organs, daily sperm production, and behavior. Toxicology and Industrial Health, 1998. 14(1-2): p. 239–260.

20. Singleton, D.W. and S.A. Khan, Xenoestrogen exposure and mechanisms of endocrine disruption. Front Biosci, 2003. 8: p. s110–s118.

21. Akingbemi, B.T., Estrogen regulation of testicular function. Reproductive Biology and Endocrinology, 2005. 3(1): p. 51.

22. Kazeto, Y., A.R. Place, and J.M. Trant, Effects of endocrine disrupting chemicals on the expression of CYP19 genes in zebrafish (Danio rerio) juveniles. Aquatic toxicology, 2004. 69(1): p. 25–34.

23. Shrader, E., et al., Proteomics in zebrafish exposed to endocrine disrupting chemicals. Ecotoxicology, 2003. 12(6): p. 485–488.

24. Hinfray, N., et al., Brain and gonadal aromatase as potential targets of endocrine disrupting chemicals in a model species, the zebrafish (Danio rerio). Environmental toxicology, 2006. 21(4): p. 332–337.

25. Mills, L.J. and C. Chichester, Review of evidence: are endocrine-disrupting chemicals in the aquatic environment impacting fish populations? Science of the Total Environment, 2005. 343(1): p. 1–34.

26. Trant, J.M., et al., Developmental expression of cytochrome P450 aromatase genes (CYP19a and CYP19b) in zebrafish fry (Danio rerio). Journal of Experimental Zoology Part A: Ecological Genetics and Physiology, 2001. 290(5): p. 475–483.

27. Kishida, M., et al., Estrogen and xenoestrogens upregulate the brain aromatase isoform (P450aromB) and perturb markers of early development in zebrafish (Danio rerio). Comparative Biochemistry and Physiology Part B: Biochemistry and Molecular Biology, 2001. 129(2): p. 261–268.

28. Menuet, A., et al., Molecular Characterization of Three Estrogen Receptor Forms in Zebrafish: Binding Characteristics, Transactivation Properties, and Tissue Distributions 1. Biology of reproduction, 2002. 66(6): p. 1881–1892.

29. Jobling, S., et al., Widespread sexual disruption in wild fish. Environmental science & technology, 1998. 32(17): p. 2498–2506.

30. Castro, L.F.C., et al., The estrogen receptor of the gastropod Nucella lapillus: modulation following exposure to an estrogenic effluent? Aquatic toxicology, 2007. 84(4): p. 465–468.

31. Ciocan, C.M., et al., Identification of reproduction-specific genes associated with maturation and estrogen exposure in a marine bivalve Mytilus edulis. PloS one, 2011. 6(7): p. e22326.

32. Sonnenschein, C. and A.M. Soto, An updated review of environmental estrogen and androgen mimics and antagonists. The Journal of steroid biochemistry and molecular biology, 1998. 65(1): p. 143–150.

33. Vandenberg, L.N., et al., Hormones and endocrine-disrupting chemicals: low-dose effects and nonmonotonic dose responses. Endocrine reviews, 2012. 33(3): p. 378–455.

34. Arukwe, A., Cellular and molecular responses to endocrine-modulators and the impact on fish reproduction. Marine Pollution Bulletin, 2001. 42(8): p. 643–655.

35. Aksglaede, L., et al., The sensitivity of the child to sex steroids: possible impact of exogenous estrogens. Human reproduction update, 2006. 12(4): p. 341–349.

36. Segner, H., Zebrafish (Danio rerio) as a model organism for investigating endocrine disruption. Comparative Biochemistry and Physiology Part C: Toxicology & Pharmacology, 2009. 149(2): p. 187–195.

37. Briggs, J.P., The zebrafish: a new model organism for integrative physiology. American Journal of Physiology-Regulatory, Integrative and Comparative Physiology, 2002. 282(1): p. R3–R9.

38. Hill, A.J., et al., Zebrafish as a model vertebrate for investigating chemical toxicity. Toxicological sciences, 2005. 86(1): p. 6–19.

39. Gu, Q., et al., Genetic ablation of solute carrier family 7a3a leads to hepatic steatosis in zebrafish during fasting. Hepatology, 2014. 60(6): p. 1929–1941.

40. Gu, Q. and Z. Cui, Putative Role of Cationic Amino Acid Transporter-3 in Murine Liver Metabolism Reply. 2015.

41. Gu, Q. and Z. Cui, Nitric Oxide as a Protector From Nonalcoholic Fatty Liver Disease Reply. 2015.

42. Gu, Q., et al., Generation and characterization of a transgenic zebrafish expressing the reverse tetracycline transactivator. Journal of Genetics and Genomics, 2013. 40(10): p. 523–531.

43. Song, G., et al., Effective gene trapping mediated by Sleeping Beauty transposon. PloS one, 2012. 7(8): p. e44123.

44. Zhai, G., et al., Sept6 is required for ciliogenesis in Kupffer’s vesicle, the pronephros, and the neural tube during early embryonic development. Molecular and cellular biology, 2014. 34(7): p. 1310–1321.

45. Yang, X., et al., Nucleoporin 62-like protein activates canonical Wnt signaling through facilitating the nuclear import of β-catenin in zebrafish. Molecular and cellular biology, 2015. 35(7): p. 1110–1124.

46. Zhong, L., et al., Investigation of effect of 17α-ethinylestradiol on vigilin expression using an isolated recombinant antibody. Aquatic toxicology, 2014. 156: p. 1–9.

47. Kimmel, C.B., et al., Stages of embryonic development of the zebrafish. Developmental dynamics, 1995. 203(3): p. 253–310.

48. Monserrat, J.M., et al., Pollution biomarkers in estuarine animals: critical review and new perspectives. Comparative Biochemistry and Physiology Part C: Toxicology & Pharmacology, 2007. 146(1): p. 221–234.

49. Heath, A.G., Water pollution and fish physiology. 1995: CRC press.

50. Scott, G.R. and K.A. Sloman, The effects of environmental pollutants on complex fish behaviour: integrating behavioural and physiological indicators of toxicity. Aquatic toxicology, 2004. 68(4): p. 369–392.

51. Medina, M.H., J.A. Correa, and C. Barata, Micro-evolution due to pollution: possible consequences for ecosystem responses to toxic stress. Chemosphere, 2007. 67(11): p. 2105–2114.

52. Wolterbeek, B., Biomonitoring of trace element air pollution: principles, possibilities and perspectives. Environmental pollution, 2002. 120(1): p. 11–21.

53. Van der Oost, R., J. Beyer, and N.P. Vermeulen, Fish bioaccumulation and biomarkers in environmental risk assessment: a review. Environmental toxicology and pharmacology, 2003. 13(2): p. 57–149.

54. Nagel, R., DarT: The embryo test with the Zebrafish Danio rerio‐‐a general model in ecotoxicology and toxicology. Altex, 2001. 19: p. 38–48.

55. Selderslaghs, I.W., et al., Development of a screening assay to identify teratogenic and embryotoxic chemicals using the zebrafish embryo. Reproductive toxicology, 2009. 28(3): p. 308–320.

56. Scholz, S., et al., The zebrafish embryo model in environmental risk assessment— applications beyond acute toxicity testing. Environmental Science and Pollution Research, 2008. 15(5): p. 394–404.

57. Sipes, N.S., S. Padilla, and T.B. Knudsen, Zebrafish—As an integrative model for twenty first century toxicity testing. Birth Defects Research Part C: Embryo Today: Reviews, 2011. 93(3): p. 256–267.

58. Padilla, S., et al., Zebrafish developmental screening of the ToxCast*™* Phase I chemical library. Reproductive Toxicology, 2012. 33(2): p. 174–187.

59. Asharani, P., et al., Toxicity of silver nanoparticles in zebrafish models. Nanotechnology, 2008. 19(25): p. 255102.

60. Oberemm, A., et al., Effects of cyanobacterial toxins and aqueous crude extracts of cyanobacteria on the development of fish and amphibians. Environmental Toxicology, 1999. 14(1): p. 77–88.

61. Johnson, A., E. Carew, and K. Sloman, The effects of copper on the morphological and functional development of zebrafish embryos. Aquatic Toxicology, 2007. 84(4): p. 431–438.

62. Lammer, E., et al., Is the fish embryo toxicity test (FET) with the zebrafish (Danio rerio) a potential alternative for the fish acute toxicity test? Comparative Biochemistry and Physiology Part C: Toxicology & Pharmacology, 2009. 149(2): p. 196–209.

63. Antkiewicz, D.S., et al., Heart malformation is an early response to TCDD in embryonic zebrafish. Toxicological Sciences, 2005. 84(2): p. 368–377.

64. van den Brandhof, E.-J. and M. Montforts, Fish embryo toxicity of carbamazepine, diclofenac and metoprolol. Ecotoxicology and environmental safety, 2010. 73(8): p. 1862–1866.

65. Jezierska, B., K. Ługowska, and M. Witeska, The effects of heavy metals on embryonic development of fish (a review). Fish Physiology and Biochemistry, 2009. 35(4): p. 625–640.

66. Midgett, M. and S. Rugonyi, Congenital heart malformations induced by hemodynamic altering surgical interventions. Frontiers in physiology, 2014. 5: p. 287.

67. Andersen, T.A., K.d.L.L. Troelsen, and L.A. Larsen, Of mice and men: molecular genetics of congenital heart disease. Cellular and Molecular Life Sciences, 2014. 71(8): p. 1327–1352.

